# Inhibition of PQS signaling by the Pf bacteriophage protein PfsE enhances viral replication in *Pseudomonas aeruginosa*

**DOI:** 10.1101/2023.08.25.554831

**Authors:** Caleb M. Schwartzkopf, Véronique L. Taylor, Marie-Christine Groleau, Dominick R. Faith, Amelia K. Schmidt, Tyrza L. Lamma, Diane M. Brooks, Eric Déziel, Karen L. Maxwell, Patrick R. Secor

## Abstract

Quorum sensing, a bacterial signaling system that coordinates group behaviors as a function of cell density, plays an important role in regulating viral (phage) defense mechanisms in bacteria. The opportunistic pathogen *Pseudomonas aeruginosa* is a model system for the study of quorum sensing. *P. aeruginosa* is also frequently infected by Pf prophages that integrate into the host chromosome. Upon induction, Pf phages suppress host quorum sensing systems; however, the physiological relevance and mechanism of suppression are unknown. Here, we identify the Pf phage protein PfsE as an inhibitor of *Pseudomonas* Quinolone Signal (PQS) quorum sensing. PfsE binds to the host protein PqsA, which is essential for the biosynthesis of the PQS signaling molecule. Inhibition of PqsA increases the replication efficiency of Pf virions when infecting a new host and when the Pf prophage switches from lysogenic replication to active virion replication. In addition to inhibiting PQS signaling, our prior work demonstrates that PfsE also binds to PilC and inhibits type IV pili extension, protecting *P. aeruginosa* from infection by type IV pili-dependent phages. Overall, this work suggests that the simultaneous inhibition of PQS signaling and type IV pili by PfsE may be a viral strategy to suppress host defenses to promote Pf replication while at the same time protecting the susceptible host from competing phages.

**Abbreviated summary:** Quorum sensing regulates phage defense in *Pseudomonas aeruginosa*. The Pf phage protein PfsE inhibits PQS-mediated quorum sensing by binding to the host enzyme PqsA, while also protecting against type IV pili-dependent phage infection. This dual inhibition strategy promotes Pf replication and safeguards the host from competing phages.

## Introduction

Quorum sensing is a cell-to-cell signaling system that allows bacteria to coordinate group behaviors (1). As bacterial populations grow, signaling molecules called autoinducers accumulate (2). At sufficiently high concentrations, autoinducers bind to and activate their cognate transcriptional regulators, allowing bacterial populations to coordinate group behaviors as a function of cell density (3).

Quorum sensing has been studied extensively in the opportunistic pathogen *Pseudomonas aeruginosa* (4). *P. aeruginosa* has three primary quorum sensing systems: Las, Rhl, and PQS. The Las and Rhl quorum sensing systems utilize the acyl-homoserine lactone autoinducer signals 3-oxo-C_12_-HSL and C_4_-HSL, respectively, while the PQS system uses the alkyl-quinolone (AQ) signals 4-hydroxy-2-heptylquinoline (HHQ) and 3,4-dihydroxy-2-heptylquinoline, also known as the *Pseudomonas* quinolone signal (PQS).

In *P. aeruginosa*, quorum sensing regulates behaviors related to biofilm formation (5) and the production of secreted virulence factors such as elastase, hydrogen cyanide, and pyocyanin (6). Quorum sensing also plays important roles in shaping the outcomes of encounters with bacteriophages through the regulation of phage defense behaviors. For example, quorum sensing downregulates expression of common cell surface receptors used by phages to infect cells (7, 8). Quorum sensing also regulates phage defense systems such as CRISPR-Cas (9, 10) and some phages encode genetic systems that are regulated by host quorum sensing and function to guide phage replication decisions (11-13).

*P. aeruginosa* strains are frequently lysogenized by filamentous Pf prophages that integrate into the bacterial chromosome (14-16). Deleting the Pf4 prophage from the *P. aeruginosa* PAO1 chromosome reduces bacterial virulence potential in mouse lung (17) and wound (18) infection models. In recent work, we made similar observations in a *Caenorhabditis elegans* nematode infection model—bacteria lacking the Pf4 prophage are less virulent compared to isogenic Pf lysogens (19). In this system, Pf4 modulates *P. aeruginosa* virulence potential by downregulating PQS signaling and reducing the production of the quorum-regulated virulence factor pyocyanin (19). However, how Pf4 suppresses PQS signaling and how PQS signaling may affect Pf4 replication is not known.

Here, we determine that the Pf4 protein PfsE (PA0721) binds to the anthranilate-coenzyme A ligase PqsA, inhibiting PQS production and thus PQS signaling. PfsE inhibition of PqsA increases Pf4 replication efficiency, consistent with a role for PQS signaling in regulating bacterial behaviors related to phage defense. Notably, PfsE has been previously characterized as an inner membrane protein that binds to the type IV pili protein PilC, which inhibits type IV pili extension and protects *P. aeruginosa* from superinfection by additional Pf4 virions or from infection by other type IV pili-dependent phages (19). We believe the simultaneous inhibition of PQS signaling and type IV pili by PfsE acts to suppress host defenses while at the same time protecting the susceptible host from competing phages.

## Results

### Pf4 replication and PQS quorum sensing are inversely regulated in *P*. *aeruginosa*

While propagating Pf4 *in vitro*, we noted that successful Pf4 infections (PAO1 + Pf4) were associated with reduced pyocyanin production by *P. aeruginosa* PAO1 (**Fig 1A and B**). Conversely, deleting the Pf4 prophage from the PAO1 chromosome (PAO1^ΔPf4^) enhances pyocyanin production (**Fig 1B and C**). Because the production of phenazines like pyocyanin is positively regulated by quorum sensing (20-22), these results suggest that Pf4 replication suppresses quorum sensing in *P. aeruginosa*. Consistently, RNA-seq revealed that numerous quorum sensing genes were significantly (false discovery rate, FDR<0.05) downregulated at least two-fold in Pf4-infected cells compared to uninfected cells (**Fig 2A and B**) (23). Accordingly, phenazine (pyocyanin) biosynthesis genes are also significantly (FDR<0.05) downregulated in Pf4-infected cells (**Fig 2C**).

**Fig 1.**
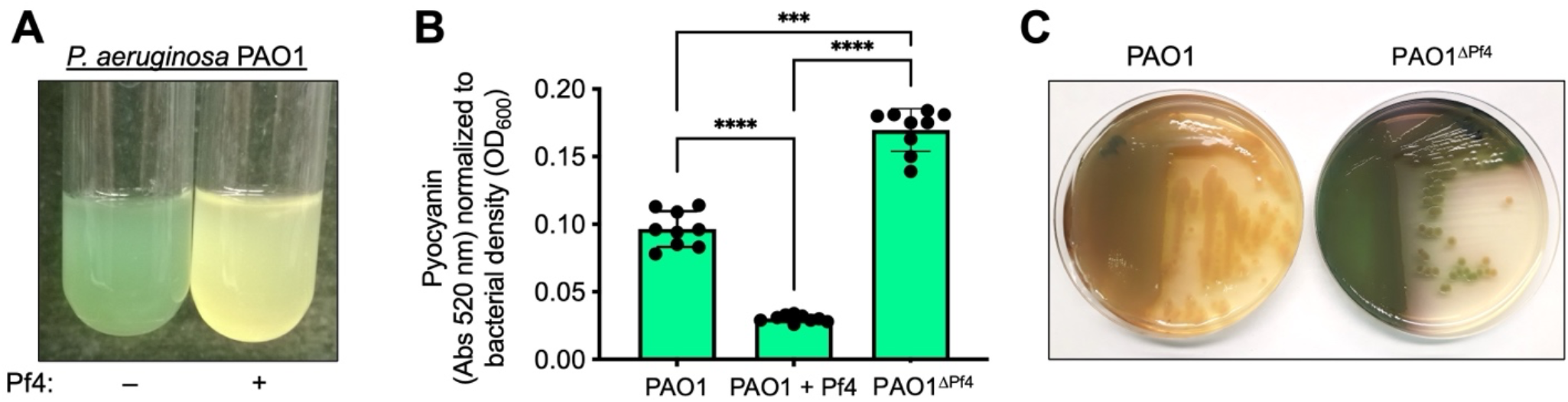
Pf4 replication and pyocyanin production are inversely regulated in *P. aeruginosa*. **(A)** Representative images of PAO1 or PAO1 superinfected with Pf4 virions (PAO1+Pf4) after 18 hours of growth in LB broth. **(B)** The green pigment pyocyanin was measured in chloroform-acid extracts of bacterial supernatants by absorbance and normalized to bacterial density (OD_600_). After 18-hours of growth, supernatants were collected from wild-type PAO1, PAO1 superinfected with Pf4 virions (PAO1 + Pf4), and PAO1 where the Pf4 prophage was deleted (PAO1^ΔPf4^). Data are the mean ±SEM of nine replicate experiments, ***P<0.001, ****P<0.0001, Student’s unpaired t-test. **(C)** Representative images of PAO1 or PAO1^ΔPf4^ grown on LB agar for 18 hours.

**Fig 2.**
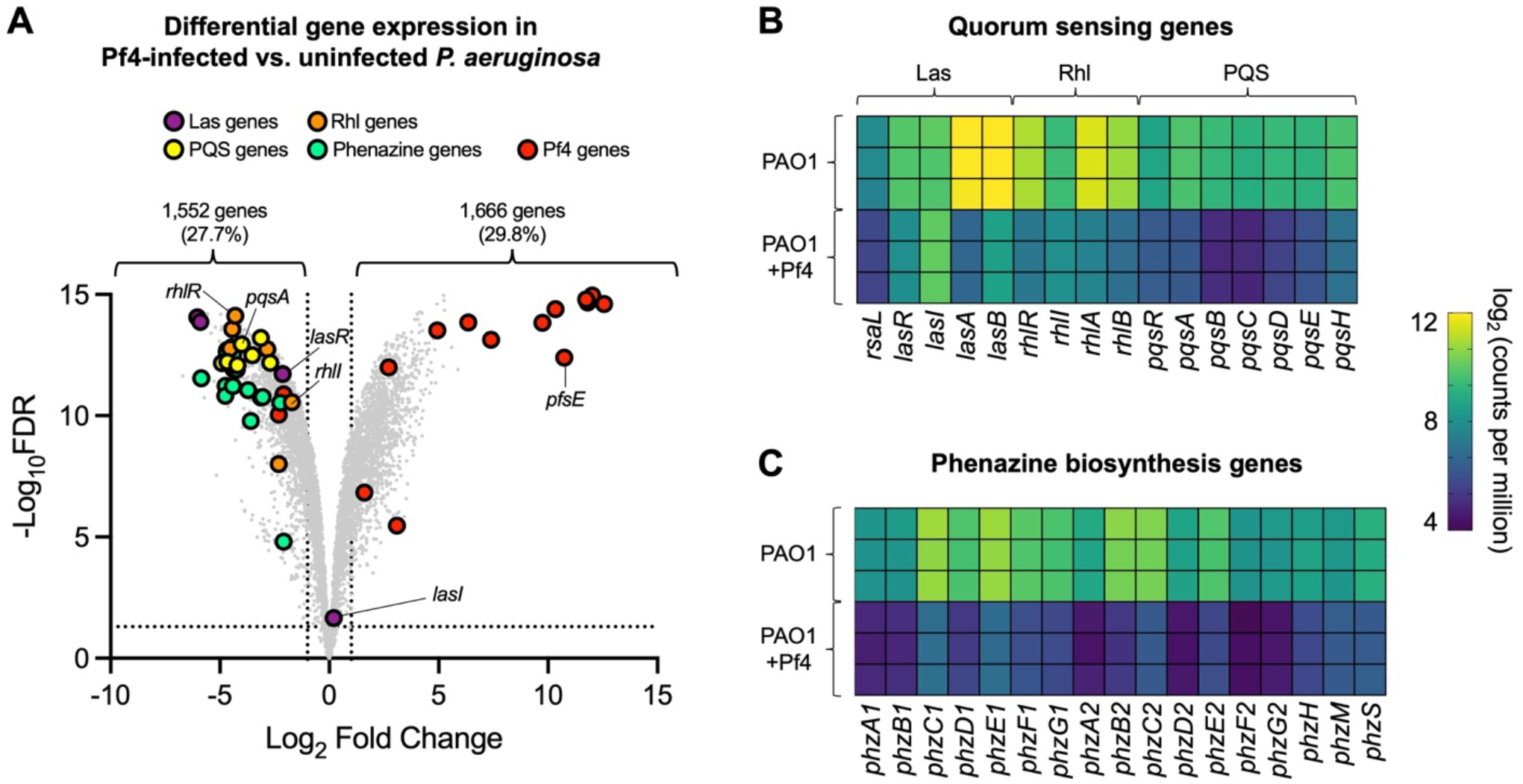
Pf4 replication downregulates *P. aeruginosa* quorum sensing and phenazine biosynthesis genes. RNAseq was performed on *P. aeruginosa* PAO1 infected with Pf4 compared to uninfected cultures in LB broth as described in reference (23). **(A)** Volcano plot showing differentially expressed genes in Pf4 infected verses uninfected *P. aeruginosa*. Dashed lines indicate differentially expressed genes that are log_2_[foldchange] > 1 and FDR<0.05 or log_2_[fold change] < -1 and FDR<0.05. Data are representative of triplicate experiments. **(B and C)** Heatmaps showing log_2_[counts per million] values for the indicated quorum sensing and phenazine biosynthesis genes are shown for each replicate.

To determine which quorum sensing systems may be affected by Pf4 infection, we used HPLC-MS and deuterated autoinducer standards to directly measure 3-oxo-C_12_-HSL, C_4_-HSL, HHQ, and PQS autoinducer levels in culture supernatants collected from PAO1, PAO1 infected by Pf4 virions (PAO1+Pf4), or PAO1^ΔPf4^ over time (**Fig 3A**). Levels of the Las autoinducer 3-oxo-C_12_-HSL were not significantly different over time in any condition (**Fig 3B**), which is consistent with the unchanged expression of the 3-oxo-C_12_-HSL autoinducer synthesis gene *lasI* (**Fig 2A and B**). Levels of the Rhl autoinducer C_4_-HSL were significantly (P<0.02) lower in Pf4-infected cells at the 12-hour time point (**Fig 3C**, red square), which is consistent with the downregulation of *rhlI* in Pf4-infected cells (**Fig 2A and B**). Collectively, these observations suggest that Pf4 replication does not drastically affect Las signaling but has a negative impact on Rhl signaling.

**Fig 3.**
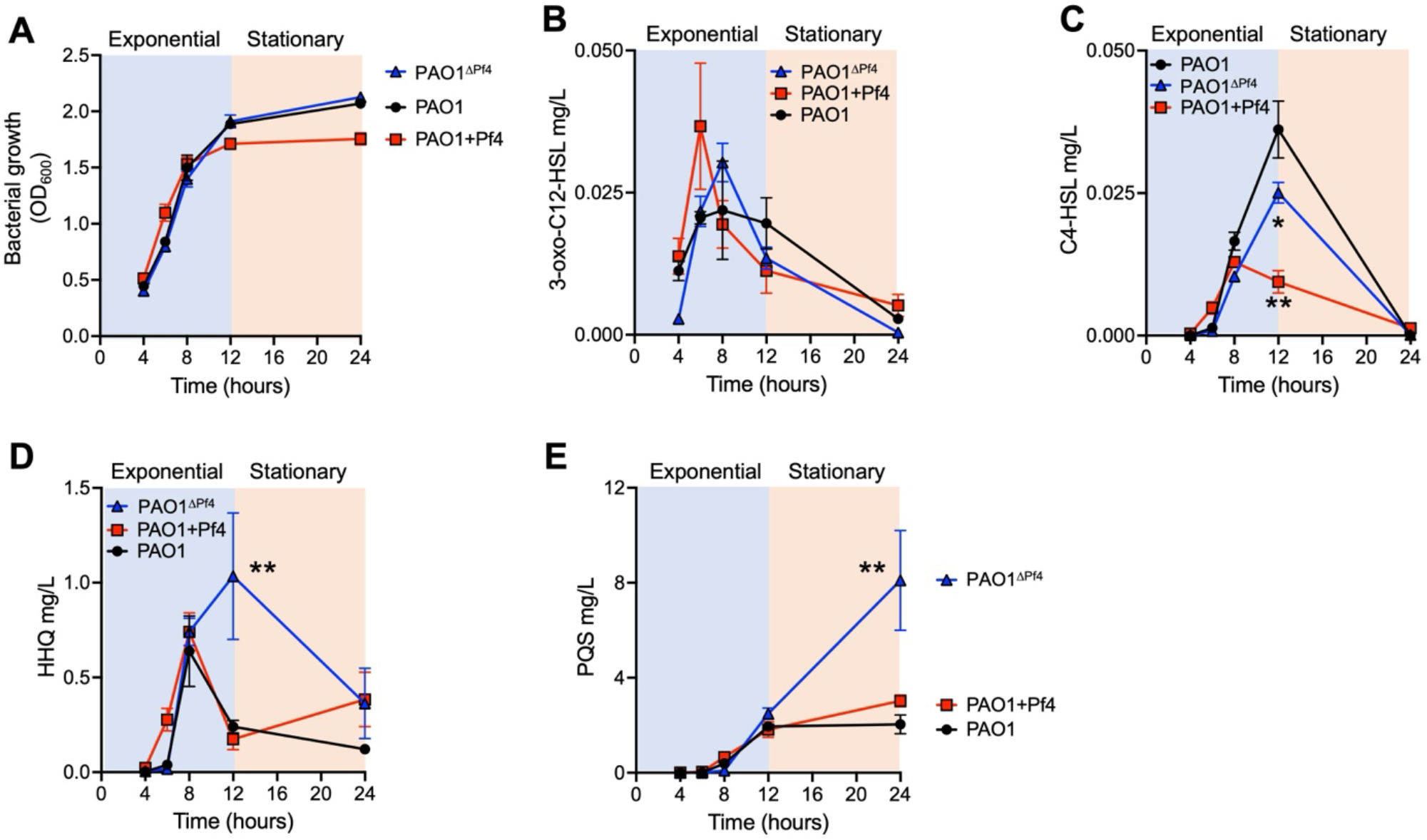
Pf4 suppresses C4-HSL, HHQ, and PQS biosynthesis. **(A)** Growth of the indicated strains was measured by absorbance at OD_600_ at the indicated times, data are the mean of three experiments. **(B-E)** The indicated quorum sensing signals in bacterial supernatants were measured by HPLC-MS at the indicated times. Data are the mean ±SEM of three replicate experiments, *P<0.05, **P<0.02 compared to PAO1 at the indicated time points.

Levels of the AQ signaling molecules HHQ and PQS were comparable in uninfected and Pf4-infected PAO1 over time (**Fig 3D** and **Fig 3E**, compare squares and circles). In the PAO1^ΔPf4^ strain, however, HHQ levels spiked at 12 hours of growth followed by a steep decline from 12 to 24 hours (**Fig 3D**, blue triangles). The decline of HHQ was accompanied by an increase in PQS levels in PAO1^ΔPf4^ supernatants from 12 to 24 hours (**Fig 3E**, blue triangles). As HHQ is the direct precursor to the PQS signaling molecule (24), these observations are consistent with HHQ being produced by PAO1^ΔPf4^ during late exponential/early stationary phase followed by HHQ conversion into PQS during stationary phase growth. These results indicate that the Pf4 prophage inhibits HHQ and PQS biosynthesis in *P. aeruginosa*.

### The Pf4 phage protein PfsE binds to PqsA

Pf4 replication suppresses the production of the quorum regulated phenazine pyocyanin (**Fig 1A**). To identify Pf4 proteins that may suppress pyocyanin production, and thus may also suppress host quorum sensing, we expressed each protein encoded by the core Pf4 genome (PA0717-PA0728) individually from an expression plasmid in *P. aeruginosa* PAO1^ΔPf4^ and measured the effects on pyocyanin production. We identified a single protein, PfsE (PA0721), that significantly (P<0.05) reduced pyocyanin production by PAO1^ΔPf4^ compared to the empty vector control (**Fig 4A**). Time course experiments confirm that expressing PfsE in PAO1^ΔPf4^ significantly (P<0.001) decreases pyocyanin production compared to PAO1^ΔPf4^ carrying an empty expression vector (**Fig 4B**). Note the *pfsE* gene is the fifth most highly upregulated gene in Pf4-infected cultures (**Fig 2A**).

**Fig 4.**
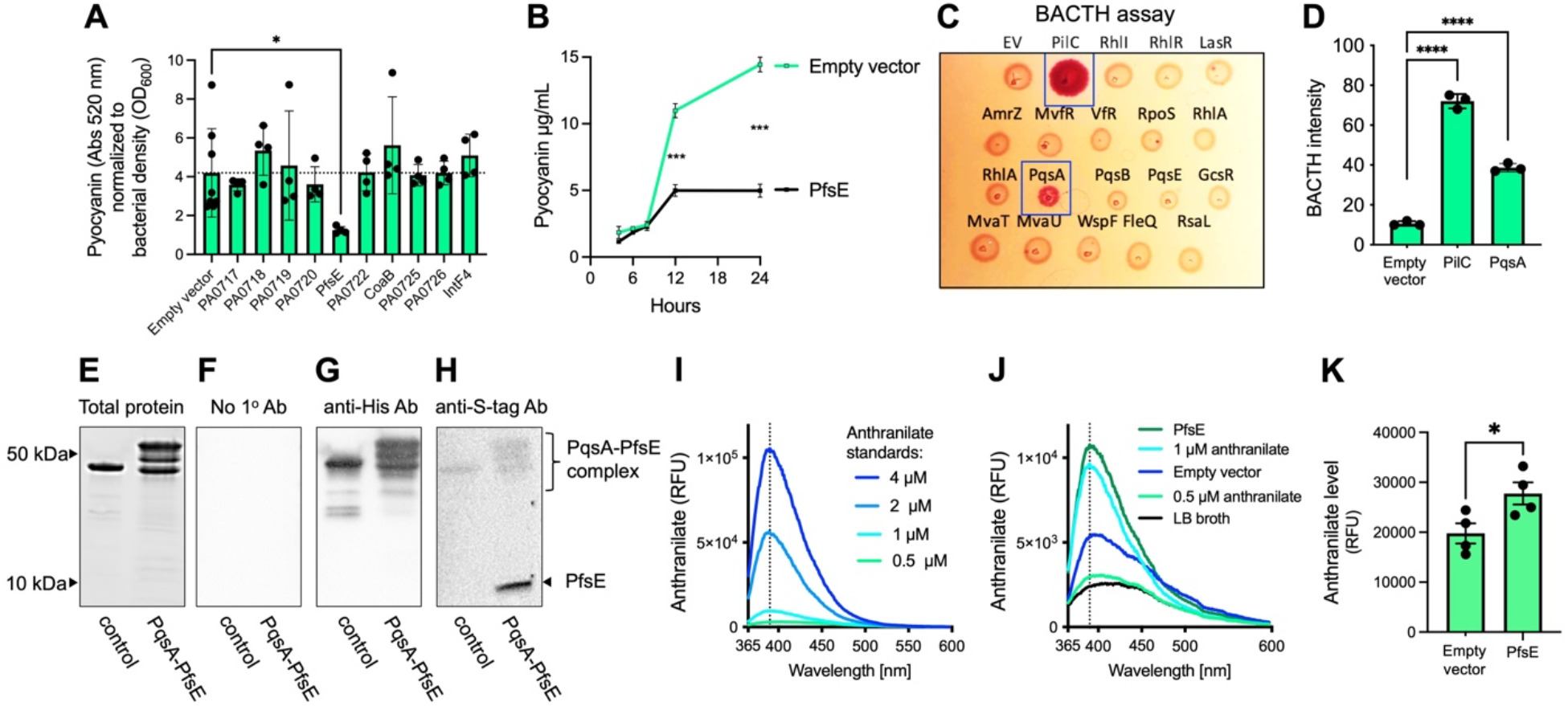
The Pf4 phage protein PfsE binds to PqsA and inhibits pyocyanin production. **(A)** The indicated Pf4 proteins were expressed from an inducible plasmid in PAO1^ΔPf4^. After 18 h, pyocyanin was extracted, quantified by absorbance, and normalized to bacterial density (OD_600_). Data are the mean +/- SEM of four experiments, *P<0.05. **(B)** Pyocyanin was extracted from PAO1^ΔPf4^ carrying an empty vector or a PfsE expression construct at the indicated times. Data are the mean +/- SEM of three experiments, ***P<0.001. **(C)** A bacterial two-hybrid assay was used to detect interactions between PfsE and the indicated bacterial proteins. Representative colonies are shown. EV = empty vector. **(D)** Pigmentation intensity of colonies expressing PfsE as bait and the indicated prey proteins was measured in image J. Data are the mean +/- SEM of three experiments, ***P<0.001. **(E-H)** His-tagged PqsA and S-tagged PfsE or His-tagged HRV-3c and S-tagged PfsE were expressed in *E. coli*. His-tagged proteins were purified from cell lysates by affinity chromatography and analyzed by SDS-PAGE and western blot using anti-His or anti-S-tag antibodies. Representative gels are shown. **(I-K)** PqsA catalyzes the conversion of anthranilate to anthraniloyl-CoA. Anthranilate levels were measured fluorimetrically in (I) LB broth spiked with anthranilate standards or (J and K) culture supernatants collected after 18 hours of growth from PAO1^ΔPf4^ carrying either an empty expression vector or PAO1^ΔPf4^ expressing PfsE. Data are the mean +/- SEM of four experiments, *P<0.03.

To determine if PfsE interacts directly with bacterial proteins involved in PQS or other quorum sensing pathways, we used a bacterial two-hybrid (BACTH) assay (25) to measure protein-protein interactions between bait (PfsE) and prey (bacterial proteins). Positive interactions are detected as red pigmentation in *E. coli* reporter colonies after 48 hours growth on MacConkey agar. PfsE is known to strongly bind the type IV pilus protein PilC (26), providing a positive control. Colony pigmentation was observed when PfsE was expressed with PilC and PqsA (**Fig 4C and D**), suggesting that in addition to PilC, PfsE also binds to PqsA.

To confirm the results from the BATCH assay, we expressed His-tagged PqsA and S-tagged PfsE in *E. coli* and purified His-tagged protein complexes by affinity chromatography (**Fig 4E**). A His-tagged HRV-3c protease (47.8 kDa) expressed in *E. coli* with PfsE was included as non-specific control. Isolated proteins were analyzed by western blot using anti-His and anti-S-tag antibodies. Blotting without a primary antibody shows no background staining (**Fig 4F**). Anti-His antibodies recognize HRV-3c and proteins isolated by affinity chromatography that range in size from ∼53–46 kDa (**Fig 4G**), indicating that His-tagged PqsA is present in the purified proteins. His-tagged PqsA has a calculated molecular weight of ∼56 kDa; however, PqsA is highly hydrophobic, which can cause it to run faster on SDS-PAGE than predicted (27), which may explain the observed reactivity in the ∼53–46 bands. The multiple bands could also be the result of PqsA proteolysis. S-tagged PfsE (3.2 kDa) was not detected in the HRV-3c sample but was detected towards the bottom of the gel in the sample containing PqsA (**Fig 4H**), suggesting that PfsE disassociates from PqsA under denaturing conditions.

PqsA catalyzes the conversion of anthranilate to anthraniloyl-CoA as a first step in HHQ and PQS biosynthesis (27). We hypothesized that PfsE binding to PqsA would cause anthranilate levels to accumulate in *P. aeruginosa* culture supernatants. To test this hypothesis, anthranilate levels were measured fluorometrically (28) in culture supernatants. Anthranilate standards are shown in **Figure 4I**. In stationary phase cells (18 h) expressing PsfE, anthranilate concentrations were approximately 1 μM (**Fig 4J**), which is ∼1.4 fold higher than supernatants collected from cells carrying an empty expression vector (**Fig 4K**). Collectively, these results suggest that PfsE binds to PqsA and inhibits its enzymatic activity.

### PfsE downregulates *pqsA* transcription

It was previously shown that Pf4 infection downregulates *pqsA* expression (23) (**Fig 2**). The expression of *pqsA* is regulated by the LysR-type regulator PqsR (also named MvfR), which binds directly to the *pqsABCDE* promoter upon binding with its cognate ligand PQS (29). We hypothesized that PfsE inhibition of PqsA and the subsequent reduction of PQS levels would downregulate *pqsA* transcription. In order to test this, we needed a mutant Pf4 prophage that lacks *psfE*. In prior work, our attempts to delete *pfsE* from the Pf4 prophage were lethal to *P. aeruginosa*; however, we were successful in deleting *pfsE* from the Pf4 integrase mutant Δ*intF4*, producing a Δ*intF/pfsE* double mutant (26). We then measured *pqsA* transcriptional reporter activity over time using a fluorescent *pqsA* transcriptional reporter (30) in PAO1 compared to PAO1^ΔPf4^ and in Δ*intF4* compared to Δ*intF4/pfsE*. We found that the transcription of *pqsA* was significantly (P<0.01) downregulated in PAO1 compared to PAO1^ΔPf4^ (**Fig 5A**), consistent with RNAseq results (**Fig 2A and B**). *pqsA* transcription was also downregulated at comparable levels in Δ*intF4* cells relative to Δ*intF4/pfsE* cells **(Fig 5A)**. Furthermore, PfsE expression significantly (P<0.01) downregulated *pqsA* transcription after 18 hours of growth in PAO1^ΔPf4^ **(Fig 5B)**. Collectively, these observations indicate that PfsE negatively regulates *pqsA* transcription.

**Fig 5.**
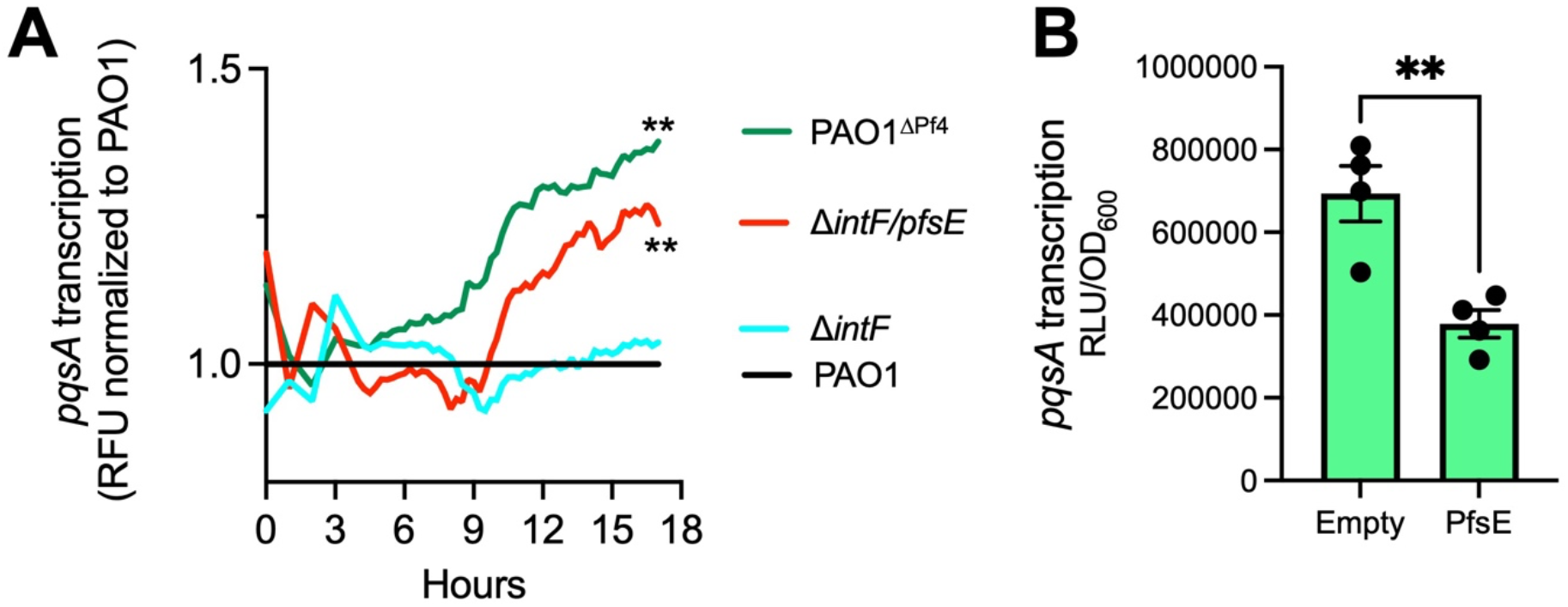
PfsE negatively regulates *pqsA* transcription. **(A)** The activity of a fluorescent P_pqsA_-*gfp* transcriptional reporter was measured in the indicated strains grown in lysogeny broth at 37°C. For each measurement, GFP fluorescence was corrected for by bacterial growth (OD_600_) and normalized to PAO1. Data are the mean of four experiments. **P<0.01, two-way ANOVA comparing PAO1 to PAO1^ΔPf4^ or Δ*intF4* to Δ*intF4/pfsE*. Error bars are omitted for clarity. **(B)** *pqsA* reporter activity was measured 18 hours post induction of expression vectors and normalized to bacterial growth (OD_600_). Data are the mean +/- SEM of four experiments, **P<0.01.

### Disabling PQS signaling enhances Pf4 replication efficiency

PqsA catalyzes the first step in PQS biosynthesis (27) and PQS signaling is implicated in regulating phage defense behaviors in *P. aeruginosa* (31-35). Thus, we hypothesized that inhibiting PQS signaling would increase *P. aeruginosa* susceptibility to Pf4 infection. To test this, we deleted *pqsA* from the PAO1^ΔPf4^ background (PAO1^ΔPf4^/Δ*pqsA*) and infected with wild-type or mutant Pf4 virions at a low multiplicity of infection (MOI) of 0.01 (one virus per 100 cells) (**Fig 6A**). Under these conditions using wild-type Pf4 virions, we did not detect any infectious virions in the supernatants of PAO1^ΔPf4^ cultures, suggesting that either PAO1^ΔPf4^ suppressed Pf4 replication or Pf4 lysogenized PAO1^ΔPf4^, converting it back into the PAO1 genotype. By contrast, infection of PAO1^ΔPf4^/Δ*pqsA* with Pf4 resulted in the production of ∼4×10^3^ PFU/mL (**Fig 6B**), showing that when PQS signaling is disabled, Pf4 replication is enhanced. When PAO1^ΔPf4^ or PAO1^ΔPf4^/Δ*pqsA* were infected with Pf4^Δ*intF4*^ virions, plaque forming units increased by several orders of magnitude to 1×10^5^ PFU/mL or 2×10^8^ PFU/mL, respectively (**Fig 6C**). Pf4^Δ*intF4*^ virions lacking the *intF4* integrase are unable to lysogenize the host and may remain in an active replication state, producing higher titers compared to wild-type Pf4. When *P. aeruginosa* PAO1^ΔPf4^ was infected with Pf4 virions lacking both the *intF4* and *pfsE* genes (Pf4^Δ*intF4/pfsE*^), phages were unable to replicate whereas Pf4^Δ*intF4/pfsE*^ virion titers were ∼4.3×10^8^ PFU/mL in the supernatants of PAO1^ΔPf4^/Δ*pqsA* cultures (**Fig 6D**). These results indicate that inhibition of PQS signaling (by PfsE or through genetic manipulation) enhances the ability of Pf4 virions to infect a naïve *P. aeruginosa* host not already infected by a Pf4 prophage.

**Figure 6.**
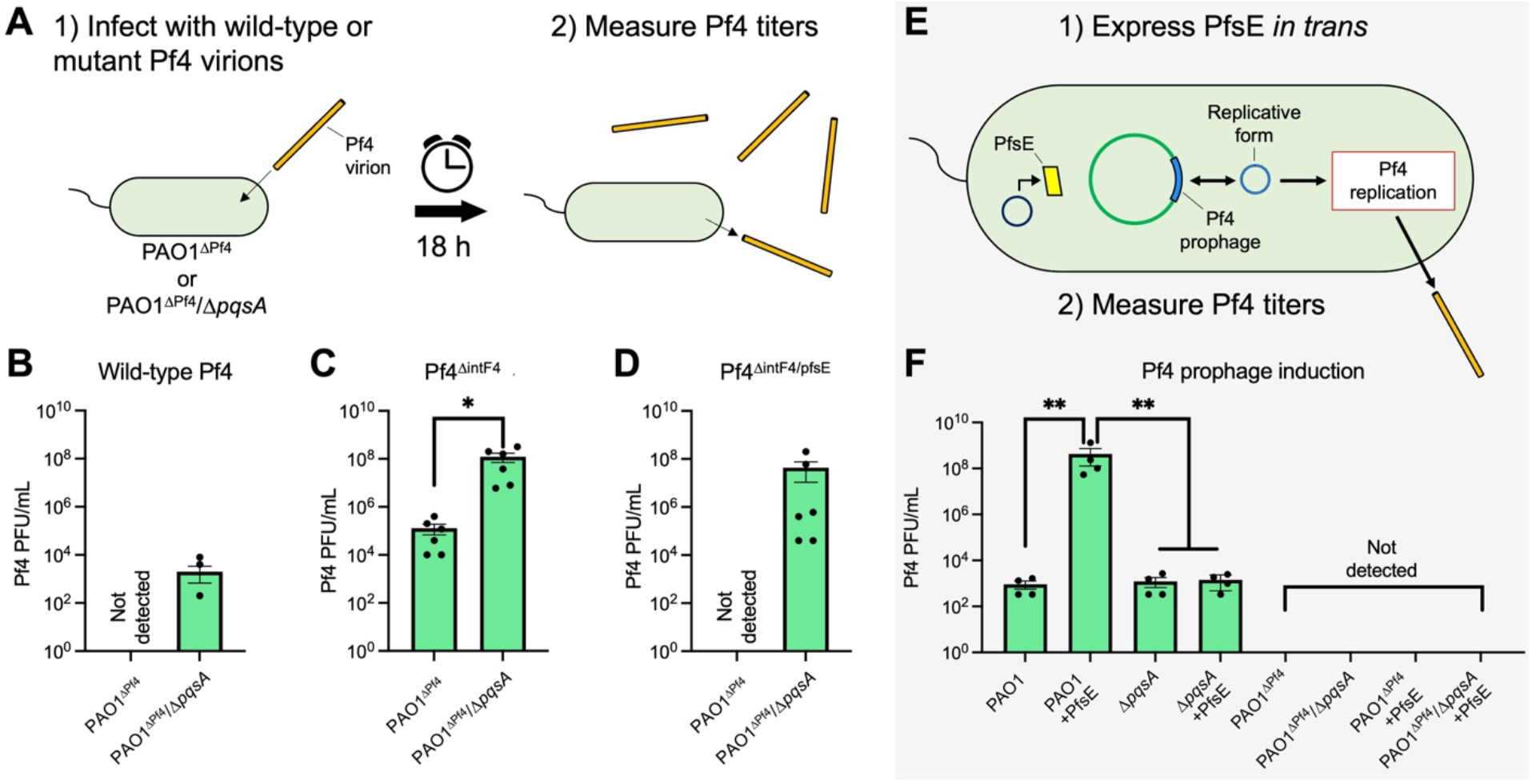
Disabling PQS signaling promotes Pf4 replication. **(A)** Liquid cultures of *P. aeruginosa* PAO1^ΔPf4^ or PAO1^ΔPf4^/Δ*pqsA* were infected with wild-type Pf4, Pf4^Δ*intF4*^, or Pf4^Δ*intF4/pfsE*^ virions (MOI 0.01). **(B-D)** After 18 hours at 37°C, Pf4 plaque forming units (PFUs) in cell culture supernatants were enumerated on lawns of *P. aeruginosa* PAO1^ΔPf4^. Data are the mean +/- SEM of 3-6 experiments, *P<0.05. **(E)** *P. aeruginosa* PAO1, PAO1^ΔPf4^, Δ*pqsA*, or PAO1^ΔPf4^/Δ*pqsA* carrying an empty expression vector or an expression vector with an inducible copy of PfsE were grown for 18 hours in LB broth at 37°C. **(F)** Pf4 PFUs in culture supernatants were then enumerated on lawns of *P. aeruginosa* PAO1^ΔPf4^. Data are the mean +/- SEM of 4 experiments, **P<0.01. Limit of detection for the assay is 333 PFU/mL.

We next tested if PQS signaling affects the transition of Pf4 from lysogeny to lytic replication. When the Pf4 prophage found in strain PAO1 is induced, it is excised from the chromosome and assumes an ∼12 kb circular double stranded DNA molecule called the replicative form (16). We hypothesized that inhibition of PQS signaling by PfsE at this critical time would be important for Pf4 to complete its lifecycle. To test this hypothesis, we expressed PfsE from an inducible plasmid in PAO1, Δ*pqsA*, PAO1^ΔPf4^, and PAO1^ΔPf4^/Δ*pqsA* and measured phage Pf4 titers in bacterial supernatants after 18 hours of growth (**Fig 6E**). In PAO1 carrying an empty expression vector, Pf4 is spontaneously produced at around 1×10^3^ PFU/mL (**Fig 6F**). In PAO1 expressing PfsE, Pf4 titers are approximetely six orders of magnitude higher at ∼1×10^9^ PFU/mL (**Fig 6F**). In Δ*pqsA* cells, Pf4 titers were comparable to those observed in PAO1 culture supernatants at ∼1×10^3^ PFU/mL and expressing PfsE in Δ*pqsA* cells did not affect Pf4 titers (**Fig 6F**), indicating inhibition of PQS signaling by PfsE is required to increase Pf4 titers. As expected, plaques were not observed under any condition where PAO1^ΔPf4^ or PAO1^ΔPf4^/Δ*pqsA* strains were used (**Fig 6F**). These results suggest that inhibition of PQS signaling by PfsE increases the spontaneous transition of Pf4 from lysogeny to active virion replication.

## Discussion

In prior work, we identified PfsE as a small, highly conserved Pf phage protein that binds to the type IV pili protein PilC to inhibit pilus extension, protecting *P. aeruginosa* from infection by competing phages (26). In this study, we characterize an additional role for PfsE—binding to PqsA and inhibiting PQS signaling (**Fig 7**). Our results indicate that inhibiting PQS signaling enhances the ability of Pf4 to infect a new host and increases Pf4 replication fidelity after the Pf4 prophage has been induced.

**Fig 7.**
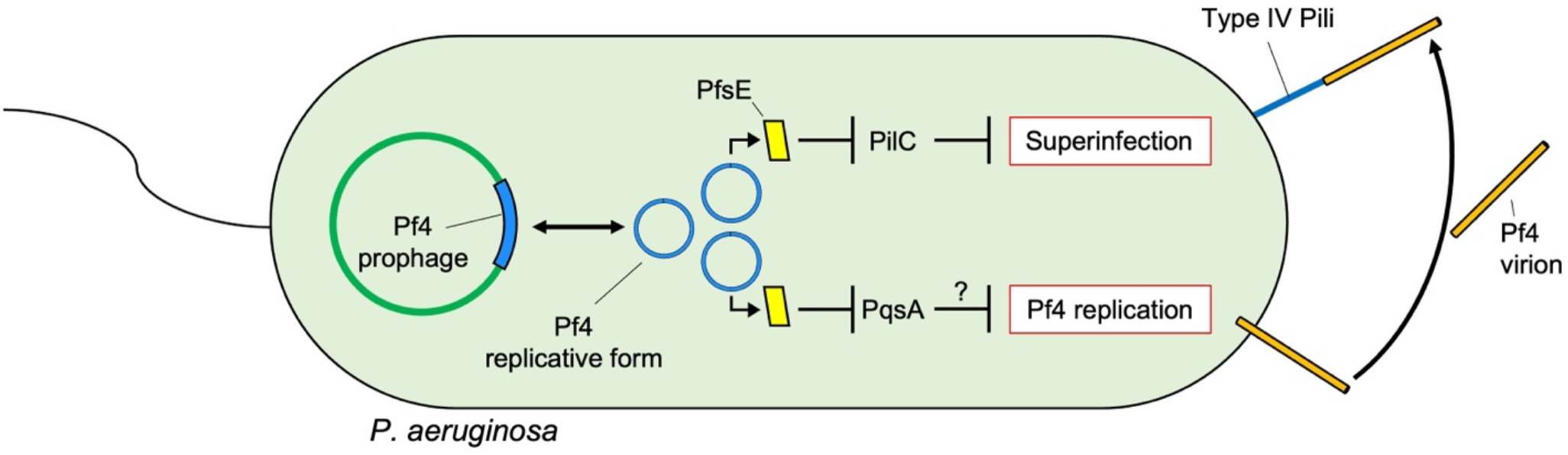
Proposed model: PfsE simultaneously inhibits PQS signaling and type IV pili extension by binding to PqsA or PilC, respectively. We propose that disabling PQS signaling promotes Pf4 replication and at the same time, protects the susceptible *P. aeruginosa* host from superinfection by Pf4 virions or infection by competing phages.

Our results are consistent with prior work indicating that PQS signaling regulates *P. aeruginosa* processes that interfere with phage replication. For example, in populations where PQS signaling is active, phage resistant isolates emerge at higher frequencies compared to populations where PQS signaling is disrupted (34). In phage infected cells, the *pqsABCDE* operon is upregulated (33) and levels of HHQ, PQS, and related metabolites are elevated (32). Furthermore, when PQS molecules are released by phage-infected cells they induce phage avoidance behavior in nearby cells (35).

*P. aeruginosa* phages have acquired mechanisms to manipulate host PQS signaling—phage JBD44 encodes genes that restore PQS signaling in quorum sensing mutants (36) while phage LUZ19 encodes a protein that binds to the PQS biosynthesis enzyme PqsD (37). These observations indicate that PQS signaling is a target in the evolutionary arms race between phages and bacteria. It is currently not known whether these other phage-encoded proteins also modulate expression of type IV pili or other cell surface receptors.

Quorum sensing regulates biofilm formation in *P. aeruginosa* (38) and Pf4 is known to be induced in *P. aeruginosa* biofilms (17, 39-44). In this study, we measured Pf4 replication primarily in liquid culture. It is possible biofilm growth could affect results. Indeed, a recent study found that Pf4 replication is induced in response to HHQ accumulation, resulting in colony biofilm autolysis (45). However, the autolysis phenotype was only observed during surface-associated growth and not in liquid culture. Our results indicate that HHQ and PQS biosynthesis is inhibited by Pf4 in liquid culture. Additional studies are required to define the relationship between PQS signaling and Pf4 replication in *P. aeruginosa* biofilms.

Our results are analogous to the Aqs1 protein encoded by the temperate *P. aeruginosa* phage DMS3. Aqs1 is a multifunctional protein that binds to and inhibits PilB and LasR to simultaneously inhibit type IV pili and disrupt *P. aeruginosa* Las quorum sensing, respectively (31). Taking our results into consideration, these observations indicate that the simultaneous inhibition of quorum sensing and type IV pili extension provides a fitness advantage to phages. However, the specific quorum sensing system that is inactivated is variable between viruses.

Overall, this study provides further insight into how Pf phages manipulate *P. aeruginosa* by inhibiting PQS signaling. Our results highlight the potential for phage-encoded proteins to influence quorum-regulated virulence and phage defense phenotypes, which has implications for therapeutic applications. For example, therapeutic phages could be engineered that encode PfsE or other phage proteins such Aqs1 that target quorum sensing pathways to simultaneously reduce pathogen virulence potential and disrupt bacterial phage defense systems, which could increase phage therapy treatment efficacy.

## Materials and methods

### Strains, plasmids, and growth conditions

Strains, plasmids, and their sources are listed in **Table 1**. Unless otherwise indicated, bacteria were grown in lysogeny broth (LB) at 37°C with 230 rpm shaking and supplemented with antibiotics (Sigma). Unless otherwise noted, gentamicin was used at 10 or 30 μg ml^−1^ and tetracycline at 100 μg ml^−1^.

**Table 1.**
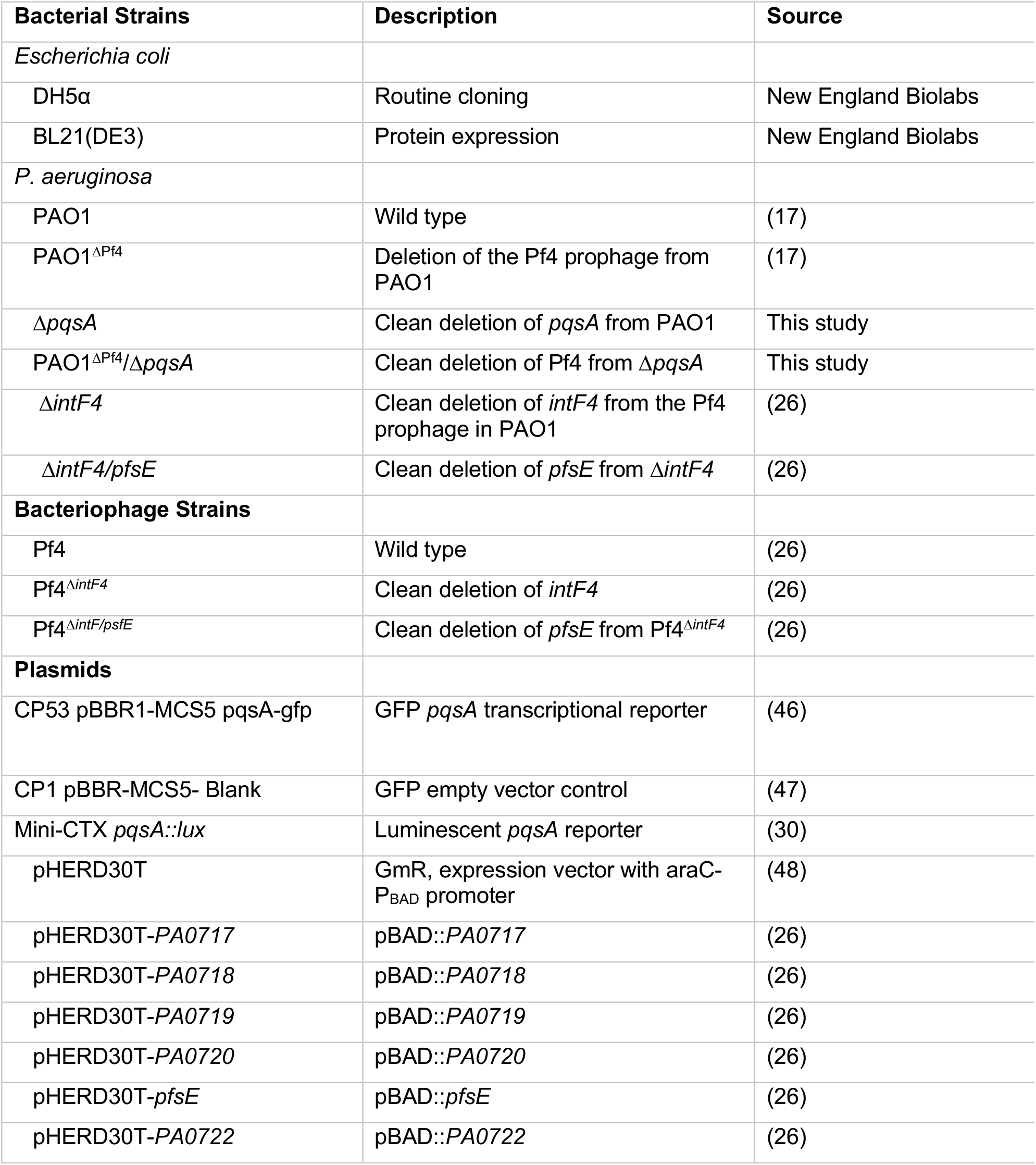
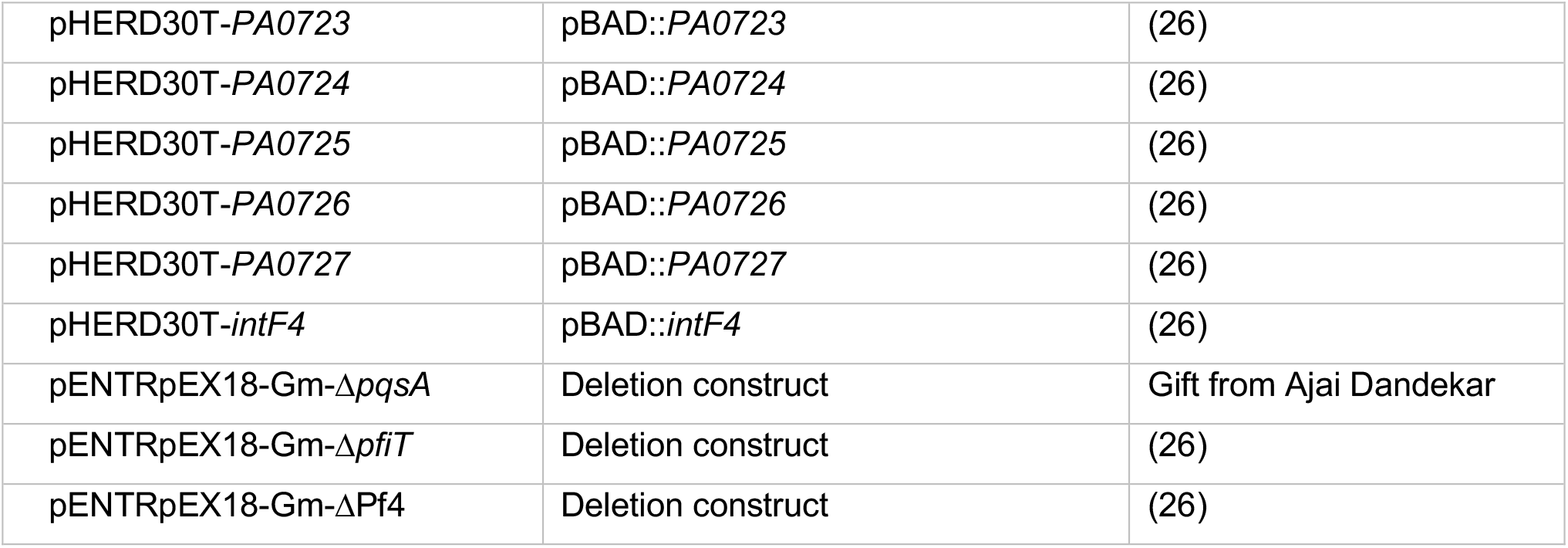
Bacterial strains, phage, and plasmids used in this study.

### Construction of *P*. *aeruginosa* mutants

All deletion strains were produced by allelic exchange (49), producing clean and unmarked deletions. All plasmids and primers used for strain construction are given in **Table 1** and **Table 2**, respectively. Briefly, using *E. coli* S17λ*pir*, we mobilized deletion constructs into *recipient strains* via biparental mating. Merodiploid *P. aeruginosa* was selected on Vogel-Bonner minimal medium (VBMM) agar containing 60 μg ml^-1^ gentamicin, followed by recovery of deletion mutants on no-salt LB (NSLB) medium containing 10% sucrose. Candidate mutants were confirmed by PCR and sequencing.

**Table 2.**
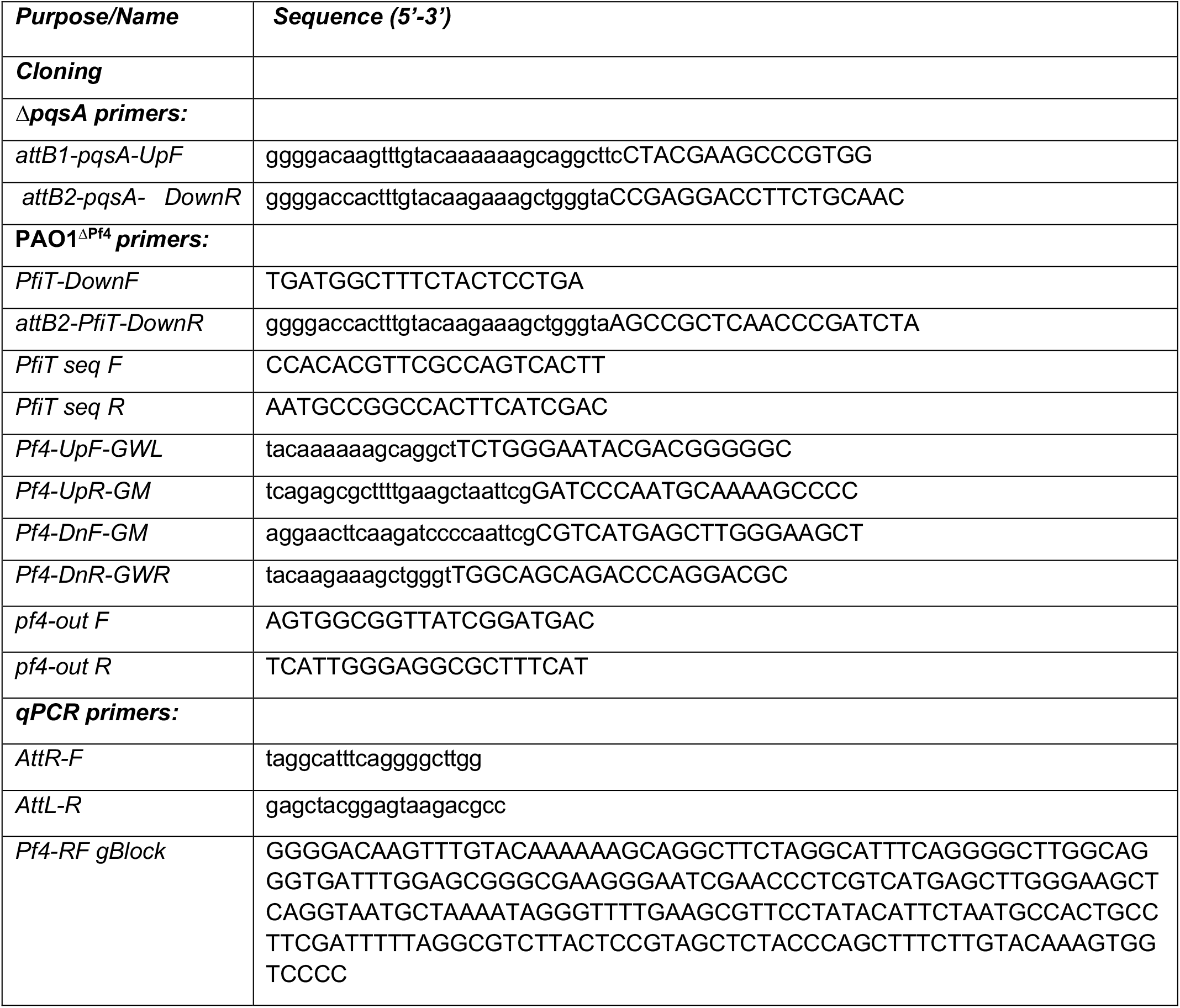
Primers used in this study. Lower case letters indicate att sites.

### Plaque assays

Plaque assays were performed using PAO1^ΔPf4^ as the indicator strain grown on LB plates. Phage in filtered supernatants were serially diluted 10x in PBS and spotted onto lawns of PAO1^ΔPf4^. Plaques were quantified after 18h of growth at 37°C.

### Pf4 phage virion enumeration by qPCR

Pf4 virion copy number was measured using qPCR as previously described (50). Briefly, filtered supernatants were treated with DNase I (10 μL of a 10mg/ml stock per mL supernatant) followed by incubation at 95°C for 10 minutes to inactivate the DNase. Ten μL reaction volumes containing 5 μl SYBR Select Master Mix (Life Technologies, Grand Island, NY), 400 nM of primer attR-F and attL-R (**Table 2**), and 1 μl supernatant. Primers attR-F and attL-R amplify the re-circularization sequence of the Pf4 replicative form (Pf4-RF) and thus, do not amplify linear Pf4 prophage sequences that may be present in contaminating chromosomal DNA. Cycling conditions were as follows: 98°C 3 min, (95°C 10 sec, 61.5°C 30 sec) x 40 cycles. A standard curve was constructed using a Pf4-RF gBlock (**Table 2**) containing the template sequence at a known copy number per mL. Pf4 copy numbers were then calculated by fitting Ct values of the unknown samples to the standard curve.

### Growth Curves

Overnight cultures were diluted to an OD_600_ of 0.01 in 96-well plates containing LB and if necessary, the appropriate antibiotics. Over the course of 24h, OD_600_ was measured in a CLARIOstar (BMG Labtech) plate reader every 15 minutes at 37°C with orbital shaking at 300 rpm for 2 minutes prior to each measurement.

### Pyocyanin extraction and quantitation

Pyocyanin was measured as described elsewhere (51, 52). Briefly, chloroform was added to culture supernatants at 50% of the total culture volume. Samples were vortexed vigorously and the different phases given time to separate (20 minutes). After the aqueous top-layer was discarded, 20% the volume of chloroform of 0.1 N HCl was added and the mixture vortexed vigorously. Once separated, the aqueous fraction was removed and absorbance at 520 nm measured. The concentration of pyocyanin in the culture supernatant, expressed as μg/ml, was obtained by multiplying the optical density at 520 nm by 17.072 (52).

### Quantification of autoinducer signalling molecules

Culture supernatants were loaded with deuterated internal standards and thrice extracted with equal parts ethyl-acetate (24). The solvent was evaporated by Savant rotorvap (Thermo RVTS10S). For quantifications of AHLs and HAQs, ethyl acetate extracts were solubilized in acetonitrile and analysed by LC/MS/MS as described (53). Briefly, samples were injected using an HPLC Waters 2795 (Mississauga, ON, Canada) on a Kinetex C18 column (Phenomenex) with an acetonitrile-water gradient containing 1% acetic acid. The detector was a tandem quadrupole mass spectrometer (Quattro premier XE; Waters) equipped with a Z-spray interface using electrospray ionization in positive mode (ESI+). Nitrogen was used as a nebulizing and drying gas at flow rates of 15 and 100 ml · min^−1^, respectively. In MRM mode the following transitions were monitored: for HHQ 244 → 159; HHQ-d_4_ 248→163; PQS 260→175; and PQS-d_4_ 264→179. The pressure of the collision gas (argon) was set at 2 × 10^−3^ mTorr and the collision energy at 30 V. For AHLs, the following transitions were monitored: C_4_-HSL 172→102; 3-oxo-C_12_-HSL 298→102 with a collision energy of 15 V.

### Anthranilate extraction and quantitation

Anthranilate was quantified as previously described (28). Briefly, fluorescence spectra (λ_ex_/λ_em_ 340 nm/365-600 nm) of sterile filtered culture supernatants were obtained on a CLARIOstar BMG LABTECH plate-reader. Standards were prepared by adding the indicated concentrations of anthranilic acid (Sigma) to sterile LB broth at room temperature.

### Quorum sensing transcriptional reporters

Competent *P. aeruginosa* PAO1, Δ*intF*, Δ*intF/pfsE*, and PAO1^ΔPf4^ were prepared by washing overnight cultures in 300 mM sucrose followed by transformation by electroporation (54) with the plasmids CP1 Blank-PBBR-MCS5 and CP53 PBBR1-MCS5 *pqsA*-gfp, listed in **Table 1**. Transformants were selected for by plating on the appropriate antibiotic selection media. The indicated strains were grown in buffered LB containing 50 mM MOPS and 100 μg ml^−1^ gentamicin for 18 hours. Cultures were then sub-cultured 1:100 into fresh LB MOPS buffer and grown to an OD_600_ of 0.3. To measure reporter fluorescence, each strain was added to a 96-well plate containing 200 μL LB MOPS with a final bacterial density of OD_600_ 0.01 and incubated at 37°C in a CLARIOstar BMG LABTECH plate-reader. Prior to each measurement, plates were shaken at 300 rpm for a duration of two minutes. A measurement was taken every 15 minutes for both growth (OD_600_) or fluorescence (excitation at 485-15 nm and emission at 535-15 nm).

pHERD30T::Empty and pHERD30T::*pfsE* were transformed into cells carrying the Mini-CTX *pqsA::lux* reporter via the same protocol as above. Strains were maintained in LB containing 100 μg ml^−1^ gentamicin and 125 μg ml^−1^ tetracycline. Cultures were then sub-cultured 1:100 into fresh LB containing 100 μg ml^−1^ gentamicin and 125 μg ml^−1^ tetracycline and grown to an OD_600_ of 0.3. To measure reporter luminescence, 200 μl aliquots were removed and luminescence was measured on a CLARIOstar BMG LABTECH plate-reader. CFU was determined using these same aliquots by 10x serial dilution and drop plating on LB agar plates with the appropriate antibiotics.

### RNA-seq data analysis

RNA-seq reads were downloaded from GEO accession no. GSE201738 (23). RNA-seq reads were then aligned to the reference *P. aeruginosa* PAO1 genome (GenBank: GCA_000006765.1), mapped to genomic features, and counted using Rsubread package v2.12.3 (55).Count tables produced with Rsubread were normalized and tested for differential expression using edgeR v3.40.2 (56).Genes with ≥ two-fold expression change and a false discovery rate (FDR) below 0.05 were considered significantly differential. RNA-seq analysis results were plotted with ggplot2 v3.4.1 and pheatmap v1.0.12 packages using R v4.2.3 in RStudio v2023.3.0.386, and GraphPad Prism v9.5.1 (57, 58).

### Anti His-tag and anti-S-tag Western blot protocol

Samples were resolved on 4-15% TGX gel. 10μg of total protein was loaded per lane. The gel was transferred to nitrocellulose and stained with Sypro Ruby (Invitrogen S11791) according to the manufacturer’s instructions. After 3 × 5minute washes with TBST (0.02M Tris Base, 0.15M NaCl, 0.05% Tween 20, pH 7.6), the membrane was blocked in TBST + 5% non-fat dry milk overnight at 4°C. The following day, the blot was washed 3 × 5minute in TBST. The blot was cut apart and the lanes with the ladders and no primary antibody were initially left in TBST. Lanes reacted with anti-His antibody were incubated for 1.5 hours in mouse anti 6X His antibody (Invitrogen MA1-21315, 1:500 in TBST) and lanes reacted with S-tag antibody were incubated in rabbit anti-mouse S-tag antibody (Invitrogen PIPA 581631, 1:500 in TBST). Blots were washed 8 × 5 minutes in TBST and then ladder lanes were reacted with Precision Plus Strep Tactin HRP Conjugate (BioRad 1610380) diluted 1:10,000 in TBST, the anti-His lanes were reacted with HRP Goat anti Mouse IgG (Abcam 6789, 1:10,000) and the S-tag lanes were reacted with HRP Goat anti Rabbit IgG (Abcam 6721, 1:25,000) for 1.5hr. The no primary antibody lanes were reacted with both HRP Goat anti Mouse and HRP Goat anti Rabbit secondary antibodies at the concentrations specified above. Blot sections were washed 5 × 5minutes in TBST and signal was detected using Clarity Western ECL Substrate (BioRad 170 5060). Images were captured on a BioRad Chemi Doc XRS+ imager.

### Statistical analyses

Unless otherwise noted, differences between data sets were evaluated with a Student’s *t*-test (unpaired, two-tailed) where appropriate. P values of < 0.05 were considered statistically significant. Area under the curve was performed 4 biological replicates. GraphPad Prism version 5.0 (GraphPad Software, San Diego, CA) was used for all analyses.

## Acknowledgments

This work was supported by NIH grants R01AI138981 and P20GM103546 to PRS. Research on the PQS system in the ED Laboratory is supported by Canadian Institutes of Health Research (CIHR) operating grant MOP-142466. The funders had no role in study design, data collection and analysis, decision to publish, or preparation of the manuscript. The authors report no conflicts of interest.

## REFERENCES

1. Miller MB, Bassler BL. 2001. Quorum sensing in bacteria. Annu Rev Microbiol 55:165–99.

2. Schuster M, Greenberg EP. 2006. A network of networks: quorum-sensing gene regulation in Pseudomonas aeruginosa. International journal of medical microbiology : IJMM 296:73–81.

3. McCready AR, Paczkowski JE, Henke BR, Bassler BL. 2019. Structural determinants driving homoserine lactone ligand selection in the Pseudomonas aeruginosa LasR quorum-sensing receptor. Proc Natl Acad Sci U S A 116:245–254.

4. Parsek MR, Greenberg EP. 2000. Acyl-homoserine lactone quorum sensing in gram-negative bacteria: a signaling mechanism involved in associations with higher organisms. Proc Natl Acad Sci U S A 97:8789–93.

5. Parsek MR, Greenberg EP. 2005. Sociomicrobiology: the connections between quorum sensing and biofilms. Trends Microbiol 13:27–33.

6. Whiteley M, Lee KM, Greenberg EP. 1999. Identification of genes controlled by quorum sensing in Pseudomonas aeruginosa. Proceedings of the National Academy of Sciences of the United States of America 96:13904–9.

7. Hoque MM, Naser IB, Bari SM, Zhu J, Mekalanos JJ, Faruque SM. 2016. Quorum Regulated Resistance of Vibrio cholerae against Environmental Bacteriophages. Sci Rep 6:37956.

8. Hoyland-Kroghsbo NM, Maerkedahl RB, Svenningsen SL. 2013. A quorum-sensing-induced bacteriophage defense mechanism. MBio 4:e00362–12.

9. Patterson AG, Jackson SA, Taylor C, Evans GB, Salmond GPC, Przybilski R, Staals RHJ, Fineran PC. 2016. Quorum Sensing Controls Adaptive Immunity through the Regulation of Multiple CRISPR-Cas Systems. Mol Cell 64:1102–1108.

10. Hoyland-Kroghsbo NM, Paczkowski J, Mukherjee S, Broniewski J, Westra E, Bondy-Denomy J, Bassler BL. 2017. Quorum sensing controls the Pseudomonas aeruginosa CRISPR-Cas adaptive immune system. Proc Natl Acad Sci U S A 114:131–135.

11. Silpe JE, Bassler BL. 2019. A Host-Produced Quorum-Sensing Autoinducer Controls a Phage Lysis-Lysogeny Decision. Cell 176:268–280 e13.

12. Silpe JE, Bassler BL. 2019. Phage-Encoded LuxR-Type Receptors Responsive to Host-Produced Bacterial Quorum-Sensing Autoinducers. MBio 10.

13. Silpe JE, Duddy OP, Johnson GE, Beggs GA, Hussain FA, Forsberg KJ, Bassler BL. 2023. Small protein modules dictate prophage fates during polylysogeny. Nature doi:10.1038/s41586-023-06376-y.

14. Knezevic P, Voet M, Lavigne R. 2015. Prevalence of Pf1-like (pro)phage genetic elements among Pseudomonas aeruginosa isolates. Virology 483:64–71.

15. Fiedoruk K, Zakrzewska M, Daniluk T, Piktel E, Chmielewska S, Bucki R. 2020. Two Lineages of Pseudomonas aeruginosa Filamentous Phages: Structural Uniformity over Integration Preferences. Genome Biol Evol 12:1765–1781.

16. Secor PR, Burgener EB, Kinnersley M, Jennings LK, Roman-Cruz V, Popescu M, Van Belleghem JD, Haddock N, Copeland C, Michaels LA, de Vries CR, Chen Q, Pourtois J, Wheeler TJ, Milla CE, Bollyky PL. 2020. Pf Bacteriophage and Their Impact on Pseudomonas Virulence, Mammalian Immunity, and Chronic Infections. Front Immunol 11:244.

17. Rice SA, Tan CH, Mikkelsen PJ, Kung V, Woo J, Tay M, Hauser A, McDougald D, Webb JS, Kjelleberg S. 2009. The biofilm life cycle and virulence of Pseudomonas aeruginosa are dependent on a filamentous prophage. The ISME journal 3:271–82.

18. Sweere JM, Van Belleghem JD, Ishak H, Bach MS, Popescu M, Sunkari V, Kaber G, Manasherob R, Suh GA, Cao X, de Vries CR, Lam DN, Marshall PL, Birukova M, Katznelson E, Lazzareschi DV, Balaji S, Keswani SG, Hawn TR, Secor PR, Bollyky PL. 2019. Bacteriophage trigger antiviral immunity and prevent clearance of bacterial infection. Science 363.

19. Schwartzkopf CM, Robinson AJ, Ellenbecker M, Faith DR, Schmidt AK, Brooks DM, Lewerke L, Voronina E, Dandekar AA, Secor PR. 2023. Tripartite interactions between filamentous Pf4 bacteriophage, Pseudomonas aeruginosa, and bacterivorous nematodes. PLoS Pathog 19:e1010925.

20. Dietrich LE, Price-Whelan A, Petersen A, Whiteley M, Newman DK. 2006. The phenazine pyocyanin is a terminal signalling factor in the quorum sensing network of Pseudomonas aeruginosa. Mol Microbiol 61:1308–21.

21. Brint JM, Ohman DE. 1995. Synthesis of multiple exoproducts in Pseudomonas aeruginosa is under the control of RhlR-RhlI, another set of regulators in strain PAO1 with homology to the autoinducer-responsive LuxR-LuxI family. Journal of bacteriology 177:7155–7163.

22. Latifi A, Winson MK, Foglino M, Bycroft BW, Stewart GS, Lazdunski A, Williams P. 1995. Multiple homologues of LuxR and LuxI control expression of virulence determinants and secondary metabolites through quorum sensing in Pseudomonas aeruginosa PAO1. Molecular microbiology 17:333–343.

23. Tortuel D, Tahrioui A, David A, Cambronel M, Nilly F, Clamens T, Maillot O, Barreau M, Feuilloley MGJ, Lesouhaitier O, Filloux A, Bouffartigues E, Cornelis P, Chevalier S. 2022. Pf4 Phage Variant Infection Reduces Virulence-Associated Traits in Pseudomonas aeruginosa. Microbiol Spectr doi:10.1128/spectrum.01548-22:e0154822.

24. Deziel E, Lepine F, Milot S, He J, Mindrinos MN, Tompkins RG, Rahme LG. 2004. Analysis of Pseudomonas aeruginosa 4-hydroxy-2-alkylquinolines (HAQs) reveals a role for 4-hydroxy-2-heptylquinoline in cell-to-cell communication. Proceedings of the National Academy of Sciences of the United States of America 101:1339–44.

25. McCallum M, Tammam S, Little DJ, Robinson H, Koo J, Shah M, Calmettes C, Moraes TF, Burrows LL, Howell PL. 2016. PilN Binding Modulates the Structure and Binding Partners of the Pseudomonas aeruginosa Type IVa Pilus Protein PilM. J Biol Chem 291:11003–15.

26. Schmidt AK, Fitzpatrick AD, Schwartzkopf CM, Faith DR, Jennings LK, Coluccio A, Hunt DJ, Michaels LA, Hargil A, Chen Q, Bollyky PL, Dorward DW, Wachter J, Rosa PA, Maxwell KL, Secor PR. 2022. A Filamentous Bacteriophage Protein Inhibits Type IV Pili To Prevent Superinfection of Pseudomonas aeruginosa. mBio doi:10.1128/mbio.02441-21:e0244121.

27. Coleman JP, Hudson LL, McKnight SL, Farrow JM, 3rd, Calfee MW, Lindsey CA, Pesci EC. 2008. Pseudomonas aeruginosa PqsA is an anthranilate-coenzyme A ligase. J Bacteriol 190:1247–55.

28. Abou-Zied OK, Al-Busaidi BY, Husband J. 2014. Solvent effect on anthranilic acid spectroscopy. J Phys Chem A 118:103–9.

29. Wade DS, Calfee MW, Rocha ER, Ling EA, Engstrom E, Coleman JP, Pesci EC. 2005. Regulation of Pseudomonas quinolone signal synthesis in Pseudomonas aeruginosa. J Bacteriol 187:4372–80.

30. Diggle SP, Matthijs S, Wright VJ, Fletcher MP, Chhabra SR, Lamont IL, Kong X, Hider RC, Cornelis P, Camara M, Williams P. 2007. The Pseudomonas aeruginosa 4-quinolone signal molecules HHQ and PQS play multifunctional roles in quorum sensing and iron entrapment. Chem Biol 14:87–96.

31. Shah M, Taylor VL, Bona D, Tsao Y, Stanley SY, Pimentel-Elardo SM, McCallum M, Bondy-Denomy J, Howell PL, Nodwell JR, Davidson AR, Moraes TF, Maxwell KL. 2021. A phage-encoded anti-activator inhibits quorum sensing in Pseudomonas aeruginosa. Mol Cell doi:10.1016/j.molcel.2020.12.011.

32. De Smet J, Zimmermann M, Kogadeeva M, Ceyssens PJ, Vermaelen W, Blasdel B, Bin Jang H, Sauer U, Lavigne R. 2016. High coverage metabolomics analysis reveals phage-specific alterations to Pseudomonas aeruginosa physiology during infection. ISME J 10:1823–35.

33. Blasdel BG, Ceyssens P-J, Chevallereau A, Debarbieux L, Lavigne R. 2018. Comparative transcriptomics reveals a conserved Bacterial Adaptive Phage Response (BAPR) to viral predation. bioRxiv doi:10.1101/248849:248849.

34. Moreau P, Diggle SP, Friman VP. 2017. Bacterial cell-to-cell signaling promotes the evolution of resistance to parasitic bacteriophages. Ecol Evol 7:1936–1941.

35. Bru JL, Rawson B, Trinh C, Whiteson K, Hoyland-Kroghsbo NM, Siryaporn A. 2019. PQS Produced by the Pseudomonas aeruginosa Stress Response Repels Swarms Away from Bacteriophage and Antibiotics. J Bacteriol 201.

36. Hoyland-Kroghsbo NM, Bassler BL. 2022. Phage Infection Restores PQS Signaling and Enhances Growth of a Pseudomonas aeruginosa lasI Quorum-Sensing Mutant. J Bacteriol 204:e0055721.

37. Hendrix H, Zimmermann-Kogadeeva M, Zimmermann M, Sauer U, De Smet J, Muchez L, Lissens M, Staes I, Voet M, Wagemans J, Ceyssens PJ, Noben JP, Aertsen A, Lavigne R. 2022. Metabolic reprogramming of Pseudomonas aeruginosa by phage-based quorum sensing modulation. Cell Rep 38:110372.

38. de Kievit TR. 2009. Quorum sensing in Pseudomonas aeruginosa biofilms. Environ Microbiol 11:279–88.

39. Secor PR, Sweere JM, Michaels LA, Malkovskiy AV, Lazzareschi D, Katznelson E, Rajadas J, Birnbaum ME, Arrigoni A, Braun KR, Evanko SP, Stevens DA, Kaminsky W, Singh PK, Parks WC, Bollyky PL. 2015. Filamentous Bacteriophage Promote Biofilm Assembly and Function. Cell Host Microbe 18:549–59.

40. Secor PR, Jennings LK, Michaels LA, Sweere JM, Singh PK, Parks WC, Bollyky PL. 2015. Biofilm assembly becomes crystal clear - filamentous bacteriophage organize the Pseudomonas aeruginosa biofilm matrix into a liquid crystal. Microb Cell 3:49–52.

41. McElroy KE, Hui JG, Woo JK, Luk AW, Webb JS, Kjelleberg S, Rice SA, Thomas T. 2014. Strain-specific parallel evolution drives short-term diversification during Pseudomonas aeruginosa biofilm formation. Proceedings of the National Academy of Sciences of the United States of America 111:E1419–27.

42. JG H, A M-p, S K, D M, SA R. 2014. Environmental cues and genes involved in establishment of the superinfective Pf4 phage of Pseudomonas aeruginosa. Front Microbiol 5.

43. Webb JS, Lau M, Kjelleberg S. 2004. Bacteriophage and phenotypic variation in Pseudomonas aeruginosa biofilm development. Journal of bacteriology 186:8066–73.

44. Whiteley M, Bangera MG, Bumgarner RE, Parsek MR, Teitzel GM, Lory S, Greenberg EP. 2001. Gene expression in Pseudomonas aeruginosa biofilms. Nature 413:860–4.

45. Giallonardi G, Letizia M, Mellini M, Frangipani E, Halliday N, Heeb S, Camara M, Visca P, Imperi F, Leoni L, Williams P, Rampioni G. 2023. Alkyl-quinolone-dependent quorum sensing controls prophage-mediated autolysis in Pseudomonas aeruginosa colony biofilms. Front Cell Infect Microbiol 13:1183681.

46. Smalley NE, Schaefer AL, Asfahl KL, Perez C, Greenberg EP, Dandekar AA. 2022. Evolution of the Quorum Sensing Regulon in Cooperating Populations of Pseudomonas aeruginosa. mBio 13:e0016122.

47. Feltner JB, Wolter DJ, Pope CE, Groleau MC, Smalley NE, Greenberg EP, Mayer-Hamblett N, Burns J, Deziel E, Hoffman LR, Dandekar AA. 2016. LasR Variant Cystic Fibrosis Isolates Reveal an Adaptable Quorum-Sensing Hierarchy in Pseudomonas aeruginosa. mBio 7.

48. Roux S, Krupovic M, Daly RA, Borges AL, Nayfach S, Schulz F, Sharrar A, Matheus Carnevali PB, Cheng JF, Ivanova NN, Bondy-Denomy J, Wrighton KC, Woyke T, Visel A, Kyrpides NC, Eloe-Fadrosh EA. 2019. Cryptic inoviruses revealed as pervasive in bacteria and archaea across Earth’s biomes. Nat Microbiol doi:10.1038/s41564-019-0510-x.

49. Hmelo LR, Borlee BR, Almblad H, Love ME, Randall TE, Tseng BS, Lin C, Irie Y, Storek KM, Yang JJ, Siehnel RJ, Howell PL, Singh PK, Tolker-Nielsen T, Parsek MR, Schweizer HP, Harrison JJ. 2015. Precision-engineering the Pseudomonas aeruginosa genome with two-step allelic exchange. Nature protocols 10:1820–41.

50. Burgener EB, Secor PR, Tracy MC, Sweere JM, Bik EM, Milla CE, Bollyky PL. 2020. Methods for Extraction and Detection of Pf Bacteriophage DNA from the Sputum of Patients with Cystic Fibrosis. Phage (New Rochelle) 1:100–108.

51. Kurachi M. 1958. Studies on the Biosynthesis of Pyocyanine.(I): On the Cultural Condition for Pyocyanine Formation. Bulletin of the Institute for Chemical Research, Kyoto University 36:163–173.

52. Essar DW, Eberly L, Hadero A, Crawford IP. 1990. Identification and characterization of genes for a second anthranilate synthase in Pseudomonas aeruginosa: interchangeability of the two anthranilate synthases and evolutionary implications. Journal of Bacteriology 172:884–900.

53. Lepine F, Milot S, Groleau MC, Deziel E. 2018. Liquid Chromatography/Mass Spectrometry (LC/MS) for the Detection and Quantification of N-Acyl-L-Homoserine Lactones (AHLs) and 4-Hydroxy-2-Alkylquinolines (HAQs). Methods Mol Biol 1673:49–59.

54. Choi K-H, Kumar A, Schweizer HP. 2006. A 10-min method for preparation of highly electrocompetent Pseudomonas aeruginosa cells: application for DNA fragment transfer between chromosomes and plasmid transformation. Journal of microbiological methods 64:391–397.

55. Liao Y, Smyth GK, Shi W. 2019. The R package Rsubread is easier, faster, cheaper and better for alignment and quantification of RNA sequencing reads. Nucleic Acids Research 47:e47–e47.

56. Robinson MD, McCarthy DJ, Smyth GK. 2009. edgeR: a Bioconductor package for differential expression analysis of digital gene expression data. Bioinformatics 26:139–140.

57. Team RC. 2014. R: A language and environment for statistical computing. MSOR connections 1.

58. Villanueva RAM, Chen ZJ. 2019. ggplot2: Elegant Graphics for Data Analysis (2nd ed.). Measurement: Interdisciplinary Research and Perspectives 17:160–167.

